# Multi-Target Mechanisms of Xiaobi Plaster in Lumbar Disc Degeneration: Insights from Network Pharmacology and Bioinformatics Analyses

**DOI:** 10.1101/2025.09.11.675695

**Authors:** Huaize Wang, Jiejie Sun, Bin Dai, Jiafeng Peng, Hongxing Zhang, Zhiwen Zheng, Minglei Gao, Ran Xu, Junchen Zhu, Yingzong Xiong

**Affiliations:** The Second Affiliated Hospital of Anhui University of Traditional Chinese Medicine, Hefei, China; Bozhou Vocational and Technical College, Bozhou, China; Graduate School, Anhui University of Traditional Chinese Medicine, Hefei, China; Department of Spine Surgery, Anhui Provincial Hospital, Hefei, China

**Keywords:** Xiaobi Plaster, Lumbar disc degeneration, Aryl hydrocarbon receptor (AHR), Network pharmacology, Molecular docking

## Abstract

**Background:** Lumbar disc degeneration (LDD) is characterized by chronic inflammation, oxidative stress, and extracellular matrix (ECM) breakdown, yet current therapies provide only symptomatic relief. Xiaobi Plaster (XBG), a traditional Chinese transdermal herbal preparation, has shown clinical benefit, but its molecular mechanisms remain unclear.

**Methods:** We applied an integrative systems biology framework combining network pharmacology, machine learning, and transcriptomic validation to identify candidate targets of Xiaobi Plaster in LDD. Molecular docking, dynamics simulations, and immune infiltration analysis further elucidated the underlying mechanisms, which were experimentally validated in a rat puncture-induced degeneration model.

**Results:** Three key targets—AHR, PTPN2, and SOAT1—were identified, with AHR emerging as the central regulator. Differential expression and enrichment analyses highlighted inflammatory and ECM-remodeling pathways, which overlapped with AHR-specific GSEA, underscoring its hub role in integrating immune and redox signaling with matrix turnover. Molecular docking and 100-ns molecular dynamics confirmed rutaecarpine (MOL002662) as the most stable AHR ligand. Transcriptomic data further showed AHR expression correlates with degeneration severity and mast cell infiltration. In vivo qPCR and Western blotting validated upregulation of AHR and supporting roles of PTPN2 and SOAT1 in degenerative discs.

**Conclusion:** This study elucidates the multi-target and multi-pathway mechanisms of XBG in LDD, highlighting AHR as a pivotal therapeutic target modulated by rutaecarpine, with PTPN2 and SOAT1 serving as auxiliary regulators. These findings provide mechanistic insights into the transdermal application of XBG and support the therapeutic relevance of targeting the AHR signaling axis in LDD.

## 1 Introduction

Lumbar disc degeneration (LDD) represents a pathological process initiated by progressive structural disruption of the intervertebral disc, which subsequently triggers abnormal cell-mediated cascades, ultimately driving extracellular matrix (ECM) degradation and functional decline [1]. Clinically, the hallmark manifestations of LDD include persistent or recurrent low back pain that may radiate to the lower extremities and, in severe cases, may be accompanied by neurological dysfunction [2]. Epidemiological studies demonstrate a striking age-related increase in prevalence: approximately 21% in individuals younger than 40 years, rising to 81% in those over 60 years, with a cumulative 25-year incidence as high as 51.5% [3]. Although topical diclofenac sodium patches, as a representative first-line pharmacological option, can provide short-term analgesic benefits, their therapeutic efficacy remains constrained by a single-target mode of action [4].

LDD is characterized by chronic inflammation, oxidative stress, ECM breakdown, and persistent pain, highlighting the need for multi-target therapeutic strategies [5,6]. Xiaobi Plaster (XBG) is a transdermal herbal preparation formulated on the basis of traditional Chinese medicine theory and accumulated clinical experience. It consists of eight herbs—Rheum palmatum (Da Huang), Scutellaria baicalensis (Huang Qin), Coptis chinensis (Huang Lian), Phellodendron amurense (Huang Bai), Notopterygium incisum (Qiang Huo), Angelica pubescens (Du Huo), Corydalis yanhusuo (Yan Husuo), and Clematis chinensis (Wei Lingxian). Historically, these herbs have been used to dispel wind-dampness, clear heat, promote blood circulation, and relieve pain in bi zheng (painful obstruction syndrome) manifesting as low back and leg pain [7–10].

The rationale for investigating XBG in LDD is further supported by evidence from related prescriptions. Duhuo Jisheng Decoction, which contains Du Huo and Qiang Huo, has been shown to alleviate disc degeneration and inhibit apoptosis in both animal models and human nucleus pulposus cells [11–13]. Similarly, Sihuang powder, which shares Huang Qin, Huang Bai, and Da Huang with XBG, significantly reduced inflammatory mediators (NO, IL-1) and attenuated cartilage degeneration in a rabbit model of synovitis [14]. Moreover, XBG itself has been evaluated clinically. A randomized study in patients with thoracic–back myofascial inflammation demonstrated that XBG significantly relieved pain and muscle tenderness, supporting its therapeutic relevance in musculoskeletal disorders [15]. In addition, modern pharmacological studies have elucidated the mechanisms of XBG’s key herbs. Yan Husuo provides potent analgesic activity through alkaloids such as tetrahydropalmatine [16]; Wei Lingxian exhibits anti-inflammatory and anti-rheumatic effects [17]; and Du Huo and Qiang Huo exert antioxidant and cartilage-protective actions [18,19]. Alongside the well-documented activities of Huang Qin [20,21], Huang Lian [22], and Huang Bo [23] in regulating inflammatory signaling and oxidative stress, these findings highlight the multi-component and multi-target nature of XBG.

Accordingly, the present study was designed to bridge this critical research gap by adopting a multidimensional methodological framework that integrates bioinformatics with artificial intelligence–driven machine learning approaches [24,25]. Specifically, we combined network pharmacology, transcriptomic enrichment analysis, and machine learning–based feature selection to systematically illuminate the multi-target molecular mechanisms of XBG at the systems level. By leveraging computational modeling alongside molecular validation, this study aims to provide robust mechanistic insights and support the clinical application of XBG in degenerative spinal conditions.

## 2 Materials and methods

### 2.1 Screening of Bioactive Components and Target Identification

This study employed a transdermal delivery-oriented strategy to identify bioactive components in XBG and their potential therapeutic targets. Using the Traditional Chinese Medicine Systems Pharmacology Database and Analysis Platform (TCMSP; http://lsp.nwu.edu.cn/tcmsp.php) [26], we screened chemical constituents from the eight herbal components of the formulation against the following ADME parameters: molecular weight (MW) ≤ 500 Da (critical threshold for skin permeation) [27], partition coefficient (logP) between 2-4 (optimizing lipid-aqueous phase balance) [28–30], and hydrogen bond donors ≤5 / acceptors ≤10 (adhering to Lipinski’s Rule of Five for passive transmembrane permeability) [30]. Compounds meeting these criteria were designated as potential transdermal bioactive agents. Target identification proceeded through a dual-path approach: First, known targets of the screened compounds were extracted directly from TCMSP. Subsequently, compound CAS numbers obtained from TCMSP were converted to SMILES structural representations via the UniProt (https://www.uniprot.org/) [31], which were then submitted to SwissTargetPrediction (http://www.swisstargetprediction.ch/) [32] to predict supplementary targets. All retrieved targets underwent standardization to official gene symbols through UniProt (Table S1).

### 2.2 Identification of Molecular Targets Associated with LDD

Targets associated with LDD were identified from transcriptomic data in the Gene Expression Omnibus (GEO) [33]. Core discovery used dataset GSE70362 (GEO accession: GSE70362) comprising 48 human intervertebral disc samples (16 non-degenerate controls, Thompson I–II; 32 LDD cases, Thompson III–V) [34]. Because our objective was to characterize disc-wide molecular changes rather than compartment-specific effects, nucleus pulposus (NP) and annulus fibrosus/fibrotic tissues were analyzed jointly to derive differentially expressed genes (DEGs) using the limma algorithm (FDR < 0.05). To verify that this joint strategy reflects shared biology across compartments, we assessed cross-tissue concordance by correlating per-gene log2 fold changes between NP and AF. Key targets were then validated in the independent dataset GSE245147 (GEO accession: GSE245147), which contains primary cultured NP cells; from this dataset we selected six untreated samples (3 non-degenerate and 3 degenerate) for expression analysis [35]. Validation included direction-of-effect consistency between discovery and validation cohorts and diagnostic performance assessment by ROC curve analysis.

### 2.3 Functional Enrichment Analysis of DEGs

Functional enrichment analysis of DEGs associated with LDD was performed through the MicrobioInfo online platform (www.bioinformatics.com.cn) for Gene Ontology (GO) and KEGG pathway assessments, applying a significance threshold of pvalue<0.05 [36]. Results were generated using a dual-display strategy: automated selection of the top five most enriched terms per category (Biological Process, Molecular Function, Cellular Component, KEGG pathways) ranked by ascending p-value, supplemented by manual curation of terms with established relevance to LDD pathogenesis. Final visualizations were rendered using ggplot2.

### 2.4 Weighted Gene Co-expression Network Analysis

This study employed the R package "WGCNA" to construct weighted gene co-expression networks for identifying functional modules coordinately regulated with the pathological progression of LDD [37]. Based on the GSE70362 gene expression dataset, preprocessed data were subjected to network construction using an optimal soft-thresholding power (scale-free topology fitting index R² > 0.8) to transform gene-gene correlation matrices into adjacency matrices, subsequently building topological overlap matrices (TOM). Hierarchical clustering with average linkage was performed using TOM-based dissimilarity measures to partition co-expressed genes into functional modules. A core module exhibiting significant positive correlation (Pearson |r| > 0.5, p< 0.05) with the LDD clinical phenotype (Thompson grades III-V) was ultimately identified, wherein genes within this module play critical roles in disease-promoting mechanisms.

### 2.5 Machine Learning-Based Feature Selection

The candidate gene set was integrated through tripartite intersection analysis of XBG drug targets, GSE70362 DEGs, and WGCNA disease-associated module genes using the R package VennDiagram. This consolidated gene set was subjected to feature selection via seven machine learning models (Random Forest, Gradient Boosting, Lasso Regression, PLS, Discriminant Analysis, Decision Tree, CatBoost), each optimized through 5-fold cross-validation repeated 10 times with gene contribution quantified by permutation importance analysis (standardized to a 0-100 scale); a dual-criterion framework was implemented for key feature identification—genes selected in ≥6 models or ranking in the top 20% by average importance score were retained as core features.

### 2.6 SHAP-Based Model Interpretability Analysis

Building upon the key feature genes identified through machine learning feature screening, a predictive modeling framework was constructed using 10 distinct algorithms: Forest, Gradient Boosting, SVM with Kernel, Logistic Model, Neighbor Method, Partial Least Squares, Neural Network, Bayes Method, Discriminant Analysis, and Lasso Regression. The model selection process employed 5-fold cross-validation repeated 10 times with receiver operating characteristic (ROC) as the primary evaluation metric. Subsequently, the optimal model determined by this procedure was retrained on the complete training set (n = 48). To deconvolute its decision mechanism, Shapley Additive exPlanations (SHAP) analysis was implemented via the R package kernelshap(v0.3.0) using 48 background samples and a custom prediction function for disease status probability output. Global feature importance ranking and directional contribution quantification were performed using the shapvizpackage (v0.9.0).

### 2.7 Cross-Dataset Expression Trend Validation

Key feature genes identified through SHAP analysis were validated for expression trend consistency across two datasets: GSE70362 (intervertebral disc tissues) and GSE245147 (nucleus pulposus cells). Group differences were visualized using box plots and diagnostic efficacy was assessed via ROC curve analysis. Expression direction consistency was rigorously examined within each dataset. Genes were excluded based on four criteria: (1) discordant expression trends across datasets, (2) single-dataset AUC < 0.55, (3) average AUC < 0.65 across both datasets, or (4) inter-dataset AUC difference > 0.20.

### 2.8 Pharmacologically Active Ingredient Retrospective Analysis

Based on the ultimately screened core genes, compound-target correspondences were extracted. Cytoscape-readable NET and ATTR attribute files were constructed and imported into Cytoscape software (v3.10.3) [38]. Three topological centrality metrics (Degree, Betweenness, and Closeness centrality) were computed using the CytoNCA plugin (v2.1.6) to analyze the systemic importance of key nodes within the network.

### 2.9 Molecular Docking Validation

After comprehensive evaluation of candidate genes across three analytical dimensions (machine learning feature importance, cross-dataset expression trends, and network topology centrality), a single core target was exclusively selected for molecular docking validation with its interacting active components. Protein crystal structures were downloaded from the PDB database [39], while compound 3D structures were obtained from the PubChem database and energy-minimized using the MMFF94 force field [40]. Molecular docking was executed with AutoDock Vina 1.2.3 [41]. Prior to docking, receptor proteins were processed in PyMOL 2.5.5 to remove water molecules, salt ions, and small molecules [42]. A docking box was then set to encompass the entire protein. Subsequently, ADFRsuite 1.0 was employed to convert all processed small molecules and receptor proteins into the PDBQT format required for AutoDock Vina 1.2.3 [43]. During docking, the global search exhaustiveness was set to 32 with other parameters maintained at default settings. The highest-scoring docking conformation was considered the binding conformation, and results were visualized using PyMOL 2.5.5 for interaction analysis.

### 2.10 Molecular dynamics simulation and binding free energy calculation

The top docking complexes were further subjected to all-atom molecular dynamics simulations using the AMBER 22 package [44]. Ligand charges were derived via the antechamber module based on Hartree–Fock SCF/6-31G* calculations performed in Gaussian 09, and subsequently parameterized with the GAFF2 force field [45–47]. Proteins were described with the ff14SB force field. Each system was solvated in a truncated octahedral TIP3P water box extending 10 Å from the solute and neutralized with counterions [48].Prior to production runs, systems underwent energy minimization, gradual heating to 298 K, and equilibration under NVT and NPT ensembles. Production MD was carried out for 100 ns at 298 K and 1 atm with a 2 fs integration step. The particle mesh Ewald method was used for long-range electrostatics, SHAKE constraints were applied to all bonds involving hydrogen atoms, and Langevin dynamics were used for temperature control. Binding free energies between proteins and ligands were estimated using the MM/GBSA approach on snapshots extracted from the final trajectories. The total free energy was decomposed into van der Waals, electrostatic, and solvation contributions, while entropy was not considered due to computational cost.

### 2.11 Clinical Correlation Analysis

The correlation between core target expression and disc degeneration severity was validated using a clinical cohort dataset with Thompson grading (Levels I-V). Thompson grades were processed as an ordered categorical variable (Level I=normal → Level V=severe degeneration); Spearman’s rank correlation test quantified the association strength; and visual representations annotated median expression trends with statistical significance.

### 2.12 Single-Gene Gene Set Enrichment Analysis (GSEA) Analysis

To dissect signaling pathways associated with the core target, single-gene GSEA was implemented: Spearman correlation coefficients between all genes and the core target were computed to generate a ranked gene list; Hallmark pathway sets (h.all.v2025.1) and Gene Ontology biological process terms (c5.go.bp.v2025.1) were loaded from the MSigDB database; pathway enrichment analysis was performed using the fgsea algorithm; significant pathways were filtered based on adjusted p-values (padj) and Normalized Enrichment Scores; and key pathways were visualized via bubble plots.

### 2.13 Immune Infiltration Analysis

To elucidate relationships between the core target and immune microenvironment, immune cell deconvolution was performed on GSE70362 intervertebral disc tissue expression data using the CIBERSORT algorithm: the LM22 signature matrix (22 immune cell subsets) calculated sample-specific immune cell proportions; group differences were assessed via Wilcoxon rank-sum test with FDR correction; immune cell interaction networks were analyzed through Spearman correlation matrices; and core target-immune cell correlations were quantified using Spearman coefficients.

### 2.14 Animal Model Construction for Disc Degeneration

The annulus fibrosus puncture method was employed to establish rat models of intervertebral disc degeneration. Thirty adult male Sprague-Dawley rats (age ≥12 weeks, weight 230–300 g) were sourced from Hangzhou Ziyuan Laboratory Animal Co., Ltd. (License: SCXK2019-0004) and housed under controlled temperature/humidity conditions with ad libitum access to food and water. Under sevoflurane anesthesia, rats were immobilized and the target disc level (coccygeal Co5/Co6 or lumbar L5/L6) was localized via fluoroscopic guidance. A 21-gauge puncture needle was percutaneously inserted perpendicularly through the full thickness of the annulus fibrosus into the nucleus pulposus center (depth 2–3 mm), rotated 180°, and maintained for 30 seconds to create controlled injury. Postoperative penicillin injections (50,000 IU/kg, intramuscular) were administered daily for 3 days to prevent infection. Disc degeneration was validated at 2 weeks post-surgery by quantifying disc height index using radiographic imaging [49]. Core target expression dynamics were assessed via Western blotting and qPCR at baseline (sham group), 1, 2, and 3 months postoperatively. All protocols were approved by the Experimental Animal Ethics Committee of the First Affiliated Hospital of USTC (Approval No.: 2023-N(A)-183).

### 2.15 Quantitative Real-Time PCR (qPCR) Analysis

In accordance with the reagent guidelines, the Trizol method (Shanghai Sangon Biotech Corporation) was utilized to extract total RNA from the spinal discs of rats. The concentration and purity of the RNA were measured using an ultra - micro - spectrophotometer (Thermo Fisher Scientific, USA), with the A260/A280 ratio ranging from 1.8 to 2.1. First - strand cDNA (Nanjing Vazyme Biotech Company, Inc., Cat. No. R122-01) was synthesized using 1 μg of RNA and the AceQ SYBR Green pre - mixed reagent (Nanjing Vazyme, Cat. No. Q111-02). The real - time quantitative PCR reaction system employed the AceQ SYBR Green pre - mixed reagent (Nanjing Vazyme, Cat. No. Q111-02) and was amplified using a LightCycler480 II fluorescence PCR instrument (Roche, Switzerland). The target genes were AHR, PTPN2, SOAT1, and GAPDH. The specific primer sequences for each gene are presented in Table S2. The procedure was as follows: pre - denaturation was carried out at 95 °C for five minutes, followed by 10 s at 95 °C and 30 s at 60 °C. The relative gene expression levels were calculated using the 2−ΔCt method. The results indicated that AHR, PTPN2, SOAT1, and GAPDH were detected by this approach.

### 2.16 Western Blot Analysis

Intervertebral discs were harvested from rat specimens, and the radioimmunoprecipitation assay (RIPA) lysis buffer was employed to extract proteins. The extracted proteins were separated via sodium dodecyl sulfate-polyacrylamide gel electrophoresis (SDS-PAGE), followed by a wet transfer onto a polyvinylidene fluoride (PVDF) membrane. To minimize non-specific binding, the membrane was incubated in a blocking buffer for one hour at room temperature. Subsequently, the membrane was incubated with primary antibodies overnight at 4 °C. The primary antibodies utilized were rabbit anti-AHR (1:2000, Proteintech, Cat No. 67785-1-Ig), rabbit anti-PTPN2 (1:5000, Proteintech, Cat No. 67388-1-Ig), rabbit anti-SOAT1 (1:1000, Proteintech, Cat No. 29369-1-AP), and rabbit GAPDH (1:50000, Proteintech, Cat No. 60004-1-Ig as an internal standard). The next day, the primary antibodies were removed, and the membrane was washed. Then, a secondary antibody conjugated with horseradish peroxidase was added and incubated for 2 h at room temperature. The chemiluminescence (CL) signal was enhanced using an enhanced chemiluminescence (ECL) reagent, and the protein signals were recorded using a CL gel imaging system (FluorChem R, Proteinsimple). The intensity of the target bands was analyzed using ImageJ software (version 1.53). Each experiment was replicated three times, and the data were normalized.

### 2.17 Statistical Analysis

Unpaired t-tests and Wilcoxon rank-sum tests were employed to assess differences between groups. Pearson correlation analysis was conducted to evaluate associations between gene expression levels. All statistical tests were two-tailed, and a p value < 0.05 was considered statistically significant. Data analysis and visualization were performed using R software (version 4.1.3).

## 3 Results

### 3.1 Screening of Potential Transdermal Bioactive Compounds and Drug Targets

According to the predefined transdermal administration-oriented ADME screening criteria, a total of 244 non-redundant potential transdermal bioactive compounds of XBP were retrieved from the TCMSP database, originating from Rhei Radix et Rhizoma (13), Scutellariae Radix (9), Coptidis Rhizoma (21), Phellodendri Cortex (56), Notopterygii Rhizoma et Radix (79), Angelicae Pubescentis Radix (56), Corydalis Rhizoma (61), and Clematidis Radix et Rhizoma (11). To systematically elucidate their pharmacological basis, a dual-path strategy was employed to obtain potential drug targets: first, known compound–target associations were directly extracted from the TCMSP database; subsequently, CAS numbers of all compounds were converted into SMILES structures and submitted to the SwissTargetPrediction platform for structure-based target prediction, thereby expanding the coverage of putative targets. After deduplication and standardization to UniProt official gene symbols, a total of 1,028 high-confidence putative targets were obtained, upon which the XBP drug–target database was constructed.

### 3.2 Identification of DEGs Associated with LDD

In order to identify the molecular targets associated with LDD, we performed a differential gene expression analysis using the GSE70362 dataset. DEGs were identified using the limma package, with a false discovery rate threshold of <0.05. This analysis resulted in the identification of 569 DEGs, of which 265 were upregulated and 304 were downregulated (Figure 1A). To further visualize the expression patterns, a heatmap was generated to display the top 20 upregulated and 20 downregulated genes (Figure 1B). To verify that the joint analysis captured disc-wide rather than compartment-specific signals, we compared per-gene log2 fold changes derived separately from nucleus pulposus and annulus fibrosus models and observed strong concordance (Spearman ρ = 0.908, p < 7.12e-224; Pearson r = 0.905; p < 1.26e−220), supporting the validity of the combined analysis (Supplementary Figure 1).

**Figure 1.**
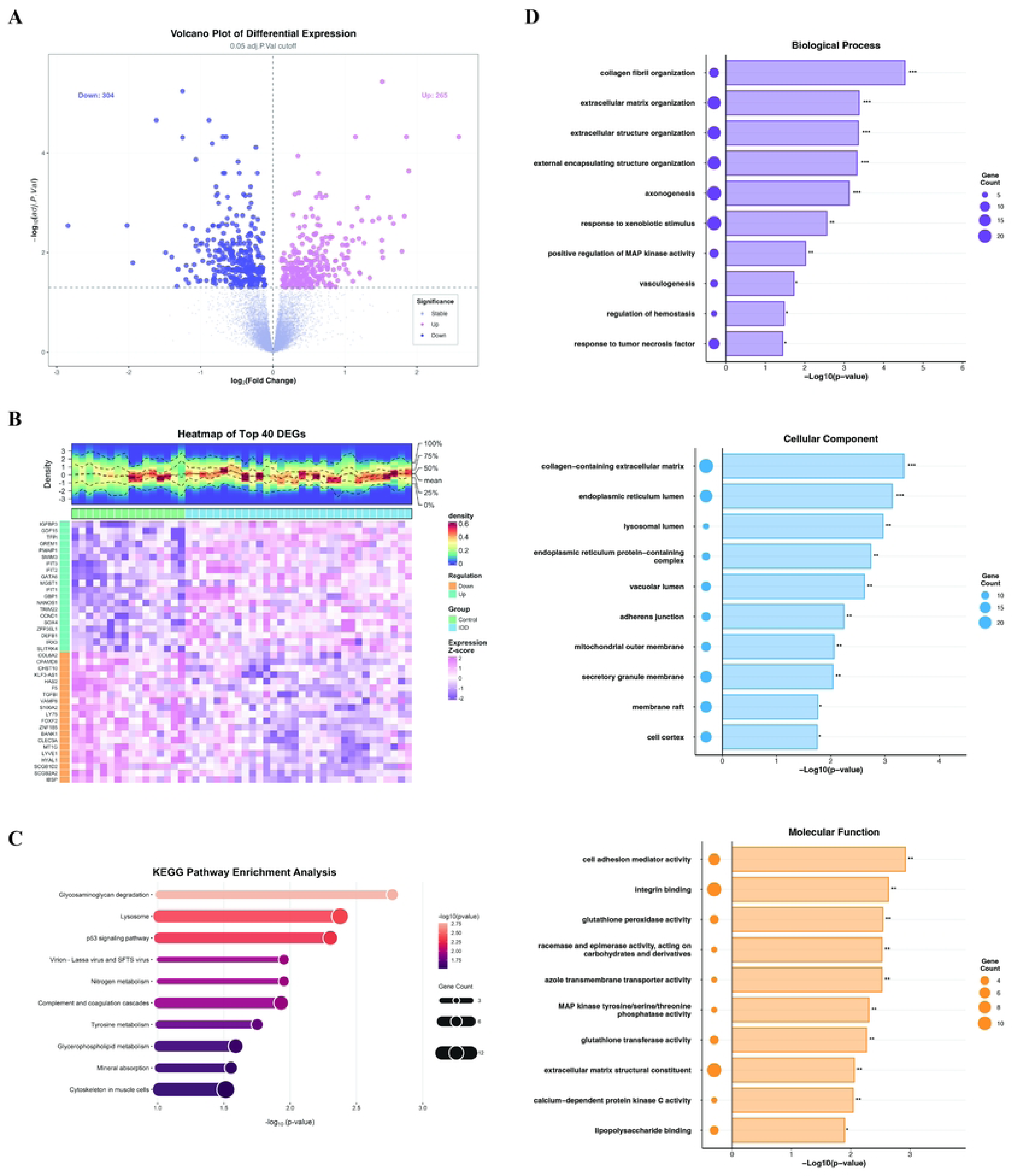
Differential expression and functional enrichment analyses in LDD. (A) Volcano plot showing upregulated (pink) and downregulated (purple) genes (FDR < 0.05). (B) Heatmap of the top 40 DEGs, illustrating distinct transcriptional patterns between degenerative and control discs. (C) KEGG pathway enrichment of DEGs, highlighting inflammation-, oxidative stress-, and ECM-related pathways. (D) GO enrichment analysis classified into Biological Process, Cellular Component, and Molecular Function categories, with terms strongly associated with ECM remodeling and stress responses. *p<0.05; **p<0.01; ***p<0.001;

### 3.3 Enrichment Analysis of DEGs

To gain insights into the biological functions and pathways associated with the DEGs, we performed Gene Ontology (GO) and Kyoto Encyclopedia of Genes and Genomes (KEGG) pathway enrichment analyses. The analysis was conducted using the MicrobioInfo online platform, with a significance threshold of p < 0.05. The GO analysis categorized the DEGs into three main categories: Biological Process, Molecular Function, and Cellular Component, (Figure 1D). In the Biological Process category, the most significantly enriched terms were associated with collagen fibril organization, ECM organization, and axonogenesis. For Molecular Function, the top enriched terms included cell adhesion mediator activity, integrin binding, and glutathione peroxidase activity. In the Cellular Component category, genes were predominantly enriched in structures such as the collagen-containing ECM and the endoplasmic reticulum lumen. Subsequently, the KEGG pathway analysis revealed that the DEGs were significantly enriched in pathways related to glycosaminoglycan degradation, lysosome function, and various signaling pathways such as the p53 and NF-kappa B signaling pathways, which are known to play crucial roles in the pathogenesis of LDD (Figure 1C).

### 3.4 Weighted Gene Co-expression Network Analysis (WGCNA)

To further investigate the molecular mechanisms underlying LDD, we performed WGCNA on the GSE70362 dataset. The sample dendrogram and associated trait information (Figure 2A) allowed us to visualize the overall structure and groupings within the dataset.The soft-thresholding power was determined by evaluating candidate thresholds from 1 to 30 using the pickSoftThreshold function (Figure 2B). The results revealed that a power value of 14 produced the highest scale-free topology fitting index (R² = 0.872). However, due to a low mean connectivity, module identification was unsuccessful. To balance the scale-free topology index (R² ≥ 0.8) and the ability to identify distinct modules, we selected a power value of 10 (R² = 0.793) to construct the co-expression network. This power value resulted in the formation of modules with clear hierarchical structures and biological relevance (Figure 2C). Subsequently, we performed a module-trait relationship analysis to identify the most relevant modules for LDD. The MEmidnightblue module, which exhibited the strongest association with LDD (Figure 2D), was selected for further investigation.Finally, to explore the relationship between module membership and gene significance, we performed a correlation analysis of genes within the MEmidnightblue module. A significant positive correlation (cor = 0.46, p = 2.2e-23) was observed (Figure 2E), suggesting that genes within this module are strongly associated with LDD and may play key roles in disease progression.

**Figure 2.**
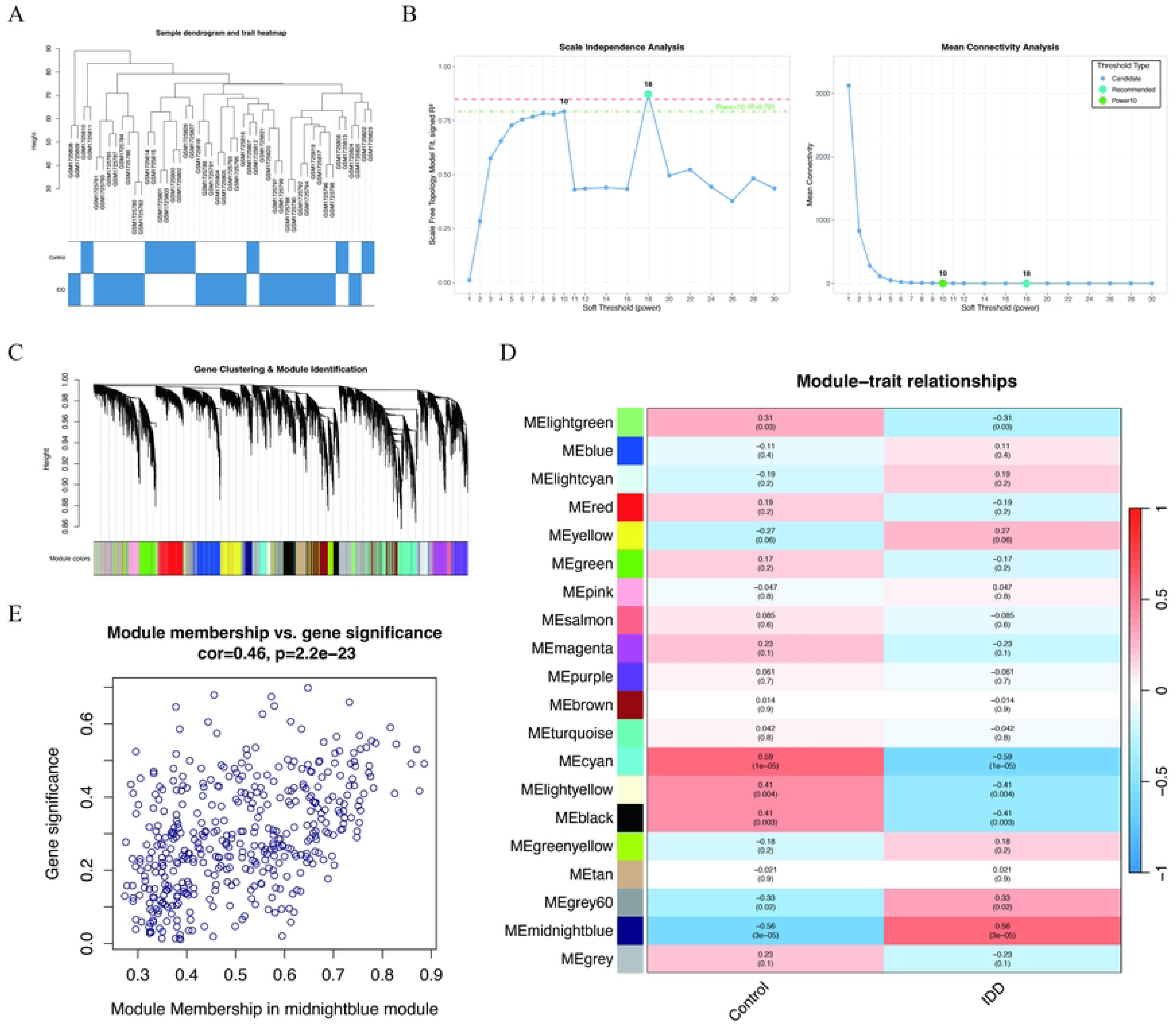
Weighted gene co-expression network analysis of LDD. (A) Sample dendrogram and trait heatmap showing clustering of samples and their association with LDD traits. (B) Scale-free topology fit index and mean connectivity analysis used to determine the optimal soft-thresholding power. (C) Gene clustering dendrogram with module detection by dynamic tree cutting, where each color represents a distinct co-expression module. (D) Module–trait correlation heatmap displaying the relationships between module eigengenes and LDD status; the midnightblue module exhibited the strongest positive correlation with degeneration. (E) Scatterplot of gene significance versus module membership in the midnightblue module, indicating a significant correlation between module connectivity and association with LDD (cor = 0.46, p = 2.2e–23).

### 3.5 Machine Learning-Based Feature Selection of Candidate Genes

To refine the list of candidate genes, a tripartite intersection analysis was performed using the VennDiagram package in R, integrating the drug target data, DEGs from the LDD dataset, and genes identified through WGCNA. This analysis revealed five overlapping genes: AHR, PTPN2, MAP3K8, SOAT1, and RB1 (Figure 3A). These genes were then subjected to feature selection using seven machine learning models—Random Forest, Gradient Boosting, Lasso Regression, PLS, Discriminant Analysis, Decision Tree, CatBoost—each optimized through 5-fold cross-validation repeated 10 times. Gene contribution was quantified using permutation importance analysis, standardized to a 0-100 scale. A dual-criterion framework was implemented to identify key features: genes selected by at least six models or those ranking in the top 20% by average importance score were retained as core features. The results of the machine learning-based feature selection are shown in (Figure 3B), where AHR consistently emerged as the most important gene across all models. This gene exhibited the highest average feature importance, confirming its significant contribution to predicting LDD. PTPN2, MAP3K8, SOAT1, and RB1 also showed notable importance but were ranked lower than AHR. In addition, the frequency with which each gene was selected by the different models is summarized in (Figure 3C). All five genes were selected by all machine learning models, indicating their consistent relevance across various algorithms.

**Figure 3.**
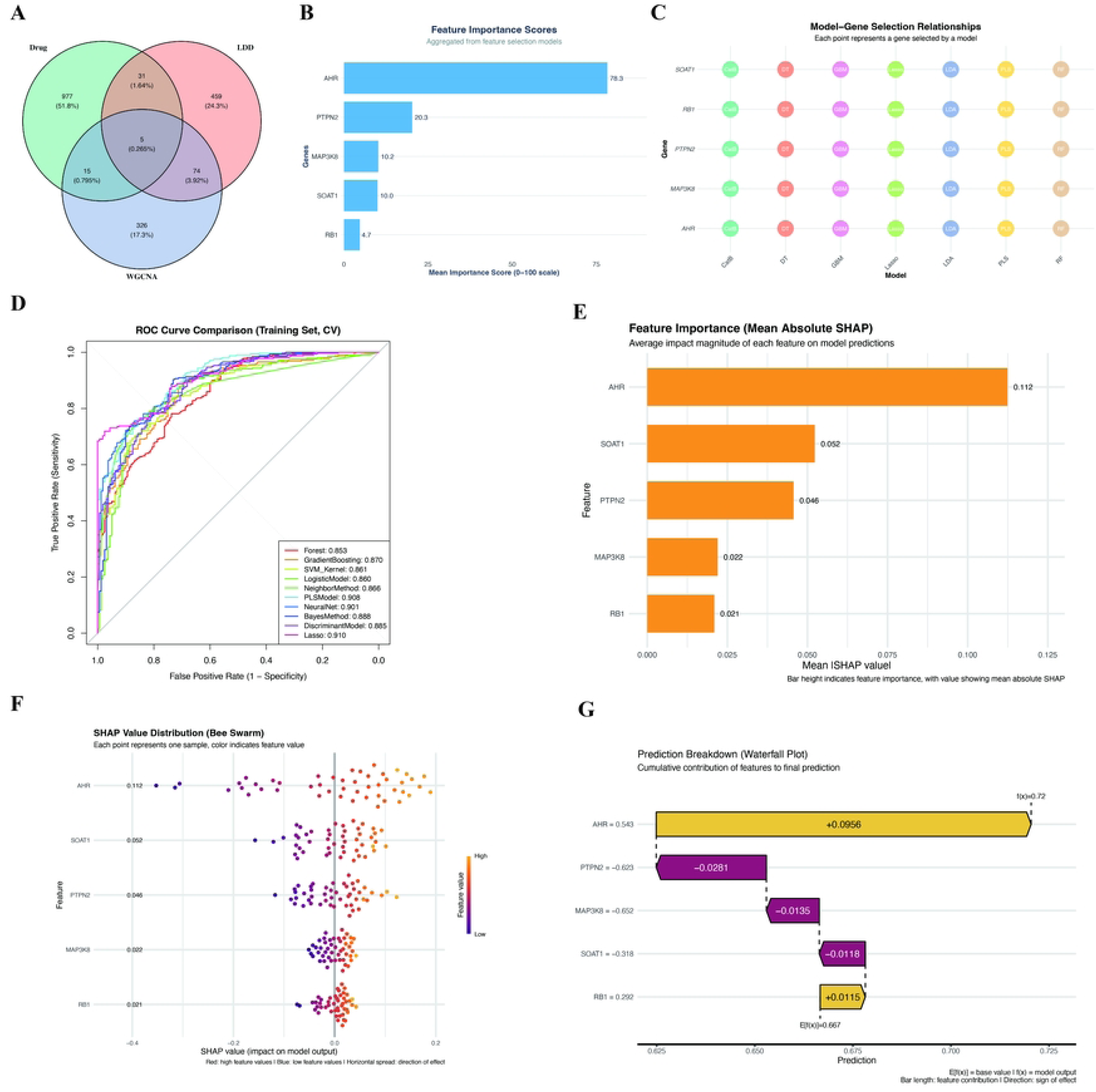
Identification and validation of hub genes through machine learning and feature selection. (A) Venn diagram showing the intersection of DEGs, WGCNA modules, and LDD-related targets. (B) Aggregated feature importance scores highlighting the top-ranked genes, with AHR emerging as the most important. (C) Model–gene selection relationships, indicating consistency across multiple algorithms. (D) ROC curve comparison of different classifiers demonstrating robust predictive performance. (E) Mean absolute SHAP values showing the relative contribution of each gene to model predictions. (F) SHAP value distribution (bee swarm plot) illustrating the direction and magnitude of each feature’s impact. (G) SHAP waterfall plot showing cumulative contributions of hub genes to an individual prediction.

### 3.6 Lasso Re-Modeling and SHAP-Based Feature Interpretation

To identify the most relevant genes for distinguishing between control and disease groups, we evaluated the performance of several machine learning algorithms, including Random Forest, Gradient Boosting, Support Vector Machine, Logistic Regression, K-Nearest Neighbors, Partial Least Squares, Neural Networks, Naive Bayes, and Lasso regression (Figure 3D). Among these, Lasso regression demonstrated the highest area under the ROC curve, indicating its superior ability to make accurate predictions. Based on this result, we decided to focus on Lasso regression for further analysis. Next, we used SHAP to interpret how each feature contributed to the model’s predictions. After re-training the Lasso regression model, SHAP analysis was applied to explain the feature importance. As shown in Figure 3E, AHR emerged as the most influential gene, exhibiting the highest mean absolute

SHAP value, followed by SOAT1, PTPN2, MAP3K8, and RB1. The lower contribution of RB1 suggests that, although it has a smaller effect in this particular model, it still plays a role. This was further confirmed by the waterfall plot (Figure 3G), which showed that RB1 had a positive contribution toward pushing the prediction towards the disease class, prompting us to retain it in the analysis, albeit with a lower priority compared to the other genes. On the other hand, MAP3K8 exhibited consistently low SHAP values across all analyses, showing minimal impact on the model’s decisions. Given this weak contribution, MAP3K8 was excluded from further analysis.

### 3.7 Validation of Expression Trends Across Independent Datasets

To validate the robustness of the identified feature genes, we examined their expression profiles in two independent cohorts. In GSE70362, AHR, PTPN2, and SOAT1 were higher in IDD than controls, whereas RB1 also showed a mild increase (Figure 4A). Single-gene ROC analysis supported good discrimination in this cohort: AHR AUC=0.904, SOAT1 AUC=0.828, RB1 AUC=0.808, and PTPN2 AUC=0.801 (Figure 4B). In GSE245147, AHR, PTPN2, and SOAT1 again increased in the IDD group, but RB1 decreased, indicating a directionally discordant trend relative to GSE70362 (Figure 4C). ROC curves reached AUC=1.000 for all four genes (Figure 4D); however, this should be interpreted cautiously given the very small sample size (3 vs 3), which increases the risk of overfitting. Taken together, AHR, PTPN2, and SOAT1 consistently demonstrated upregulation across datasets and were retained as robust candidate biomarkers, whereas RB1 showed an opposite expression pattern and was therefore excluded from further consideration.

**Figure 4.**
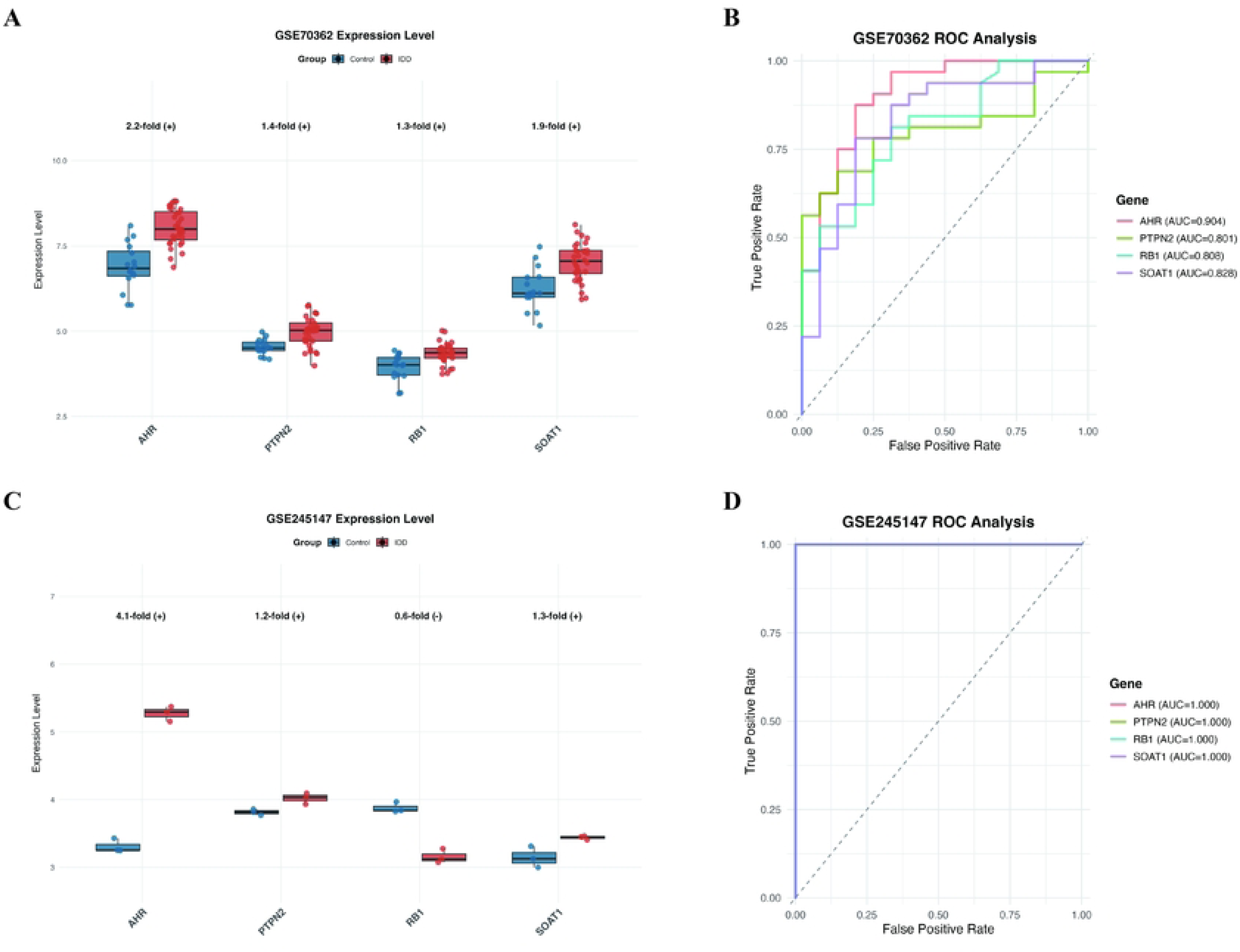
Validation of hub gene expression and diagnostic performance in independent datasets. (A) Expression levels of AHR, PTPN2, RB1, and SOAT1 in the GSE70362 dataset, showing significant upregulation of AHR, PTPN2, and SOAT1, while RB1 was downregulated in LDD compared with controls. (B) ROC curve analysis of the same genes in GSE70362, demonstrating robust diagnostic value, particularly for AHR (AUC = 0.904). (C) Validation of gene expression patterns in the independent GSE245147 dataset, confirming consistent upregulation of AHR and SOAT1, modest increase in PTPN2, and reduced RB1 expression in LDD. (D) ROC analysis in GSE245147 further supported excellent predictive performance for all four genes.

### 3.8 Compound–Gene Interaction Network Analysis

To further explore the potential pharmacological mechanisms, candidate compounds targeting the three validated genes (AHR, PTPN2, and SOAT1) were collected and used to construct a compound–gene interaction network (Figure 5A). The network was imported into Cytoscape and analyzed using the cytoNCA plugin to evaluate topological parameters, including degree, betweenness, and closeness centrality.The results highlighted PTPN2 as the central hub, exhibiting the highest degree (20) and betweenness (895.0), followed by AHR (degree = 16, betweenness = 754.0) and SOAT1 (degree = 5, betweenness = 219.0). These metrics indicate that PTPN2 and AHR occupy key regulatory positions within the compound–gene interaction network, suggesting that pharmacological modulation of these targets may exert broader biological effects.

**Figure 5.**
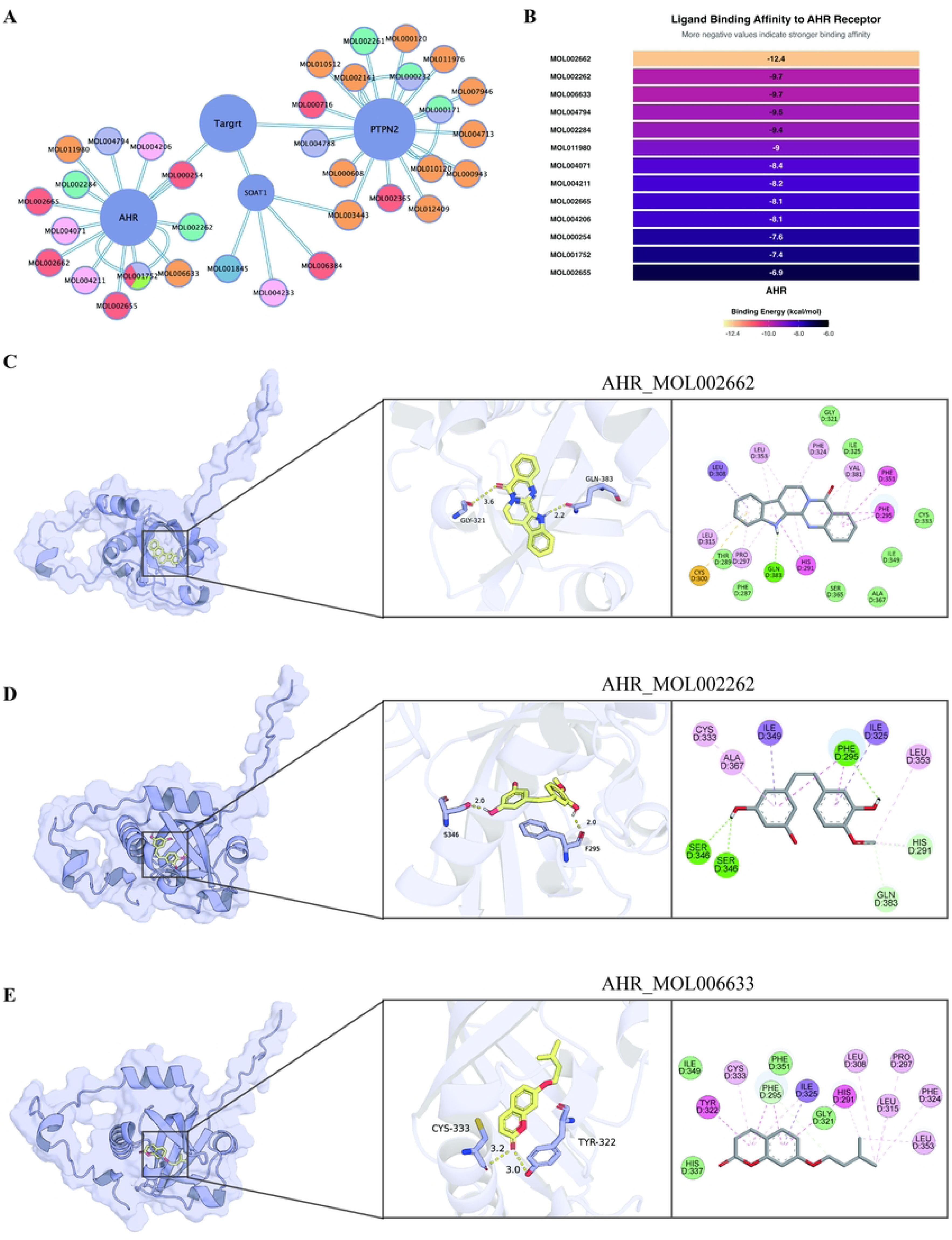
Compound–target interaction network and molecular docking of Xiaobi Plaster (XBP) bioactive components with AHR. (A) Compound–gene interaction network of Xiaobi Plaster showing predicted associations between active compounds and hub targets (AHR, PTPN2, SOAT1). (B) Binding affinities of candidate ligands to the AHR receptor, represented as docking scores (kcal/mol). All compounds showed binding energies < −5 kcal/mol, indicating favorable affinity, with rutaecarpine (MOL002662) exhibiting the strongest binding. (C–E) Representative docking poses and interaction diagrams of AHR with MOL002662 (C), MOL002262 (D), and MOL006633 (E).

### 3.9 Molecular docking validation of the core target

Given its central role across machine learning prioritization, cross-dataset expression validation, and network pharmacology topology, AHR was ultimately identified as the core target for molecular docking validation. All compounds associated with AHR were subjected to docking, and every ligand achieved a binding energy below −5 kcal/mol, indicating favorable binding affinity (Figure 5B) [50]. Among them, MOL002662 exhibited the strongest binding (−12.4 kcal/mol), followed by MOL002262 (−9.7 kcal/mol) and MOL006633 (−9.6 kcal/mol). Detailed visualization of the top three docking poses demonstrated their stable and specific interactions within the AHR binding pocket (Figure 5C–E). Collectively, these docking results confirm the structural feasibility of AHR–ligand binding and reinforce AHR as a promising therapeutic target for LDD.

### 3.10 Molecular docking validation of the core target

To further evaluate the stability and binding characteristics of the AHR–ligand complexes, 100 ns all-atom molecular dynamics simulations were conducted. As shown in (Figure 6A), the RMSD trajectories of all three complexes stabilized after ∼20 ns, with fluctuations remaining within 3–5 Å, indicating overall conformational stability. The radius of gyration (Figure 6B) remained stable throughout the simulation, supporting the compactness of the protein–ligand complexes. Ligand RMSD values (Figure 6C) confirmed that all compounds maintained relatively stable binding poses, with MOL002662 showing the lowest deviation, suggesting its optimal fit within the AHR binding pocket. RMSF analysis (Figure 6D) revealed that most residues displayed low flexibility, with moderate fluctuations restricted to loop regions. Hydrogen bond analysis (Figure 6E) showed that MOL002662 maintained the highest and most persistent number of hydrogen bonds across the trajectory, followed by MOL002262 and MOL006633, highlighting its stronger interaction stability. Binding free energy decomposition using the MM/GBSA method (Figure 6F) demonstrated consistently favorable ΔG values for all three complexes, with MOL002662 exhibiting the lowest free energy. The dominant contributions arose from van der Waals and electrostatic interactions, confirming the strong binding affinity of MOL002662. Finally, the free energy landscape (Figure 6G) revealed that the AHR–MOL002662 complex occupied a well-defined low-energy basin, corresponding to a stable conformational state with minimal structural fluctuations.

**Figure 6.**
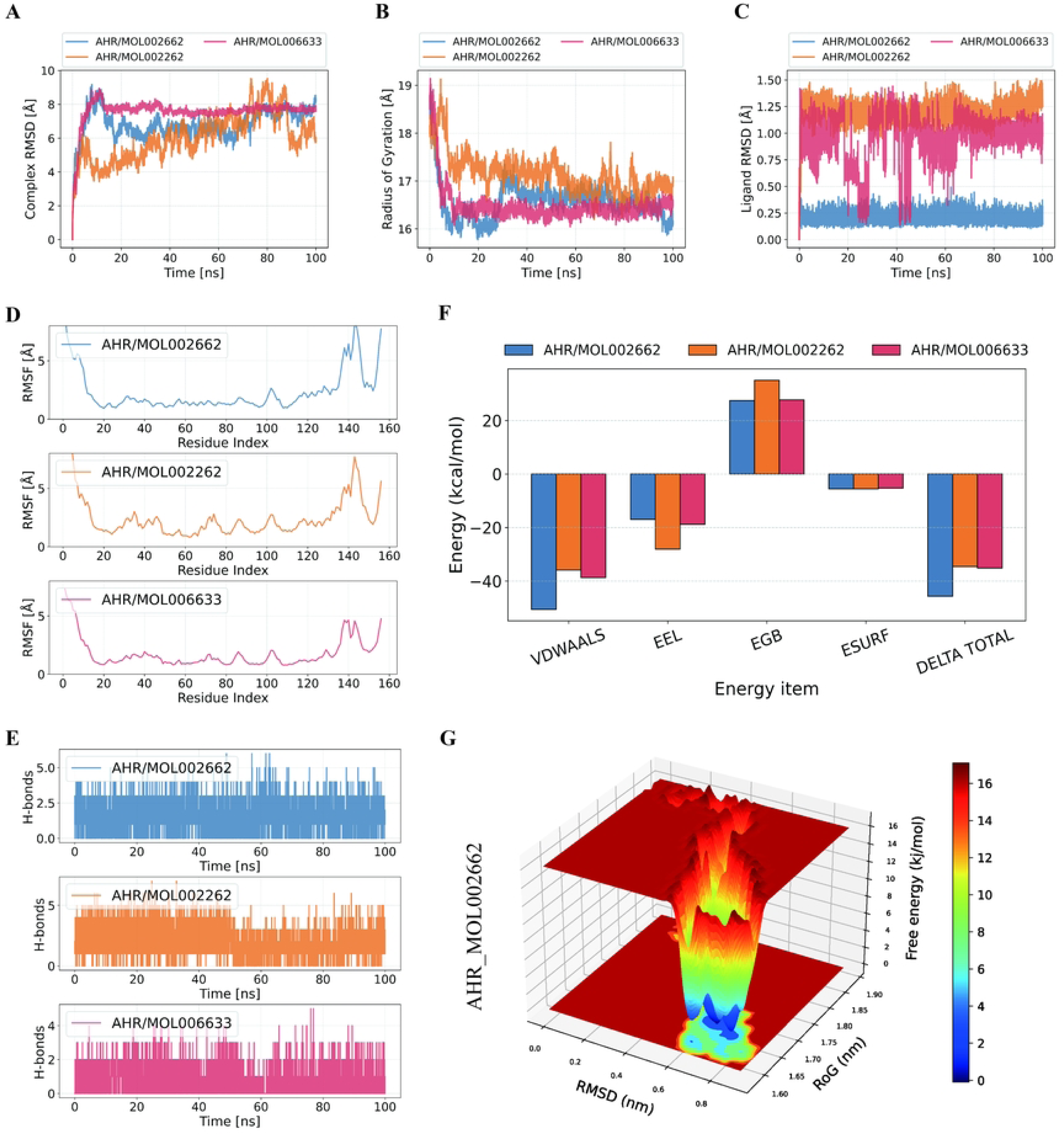
Molecular dynamics simulations of AHR–ligand complexes. (A) Time-dependent root mean square deviation (RMSD) of protein–ligand complexes, showing overall structural stability across 100 ns trajectories. (B) Radius of gyration (Rg) indicating compactness of the complexes. (C) Ligand RMSD values reflecting positional fluctuations within the AHR binding pocket. (D) Root mean square fluctuation (RMSF) of residues for AHR in complex with MOL002662, MOL002262, and MOL006633. (E) Number of hydrogen bonds formed between AHR and each ligand during the simulation period. (F) MM-PBSA binding free energy decomposition of van der Waals (VDWAALS), electrostatic (EEL), electrostatic solvation (EGB), and non-polar solvation (ESURF) components, with total ΔG values indicating binding affinity. (G) Free energy landscape (FEL) of the AHR– MOL002662 complex plotted against RMSD and Rg, highlighting the lowest-energy conformational states.

Taken together, the combined evidence from molecular docking and molecular dynamics analyses highlighted MOL002662 as the most promising and stable compound targeting AHR.

### 3.11 Clinical Correlation of AHR Expression with Thompson Grading

To further investigate the clinical relevance of the identified core gene, we analyzed AHR expression across different Thompson grades of intervertebral disc degeneration (Figure 7A). AHR expression exhibited a slight decline from grade I to grade II, followed by a progressive increase from grade III onward, reaching its highest level in advanced degeneration (grades IV–V). Correlation analysis revealed a significant positive association between AHR expression and Thompson grade (Spearman’s ρ = 0.575, p < 0.001), indicating that AHR levels generally rise with the severity of disc degeneration. The initial decline of AHR expression in early degeneration may reflect compensatory responses aimed at maintaining disc homeostasis, whereas its subsequent elevation in advanced grades indicates sustained activation during disease progression.

**Figure 7.**
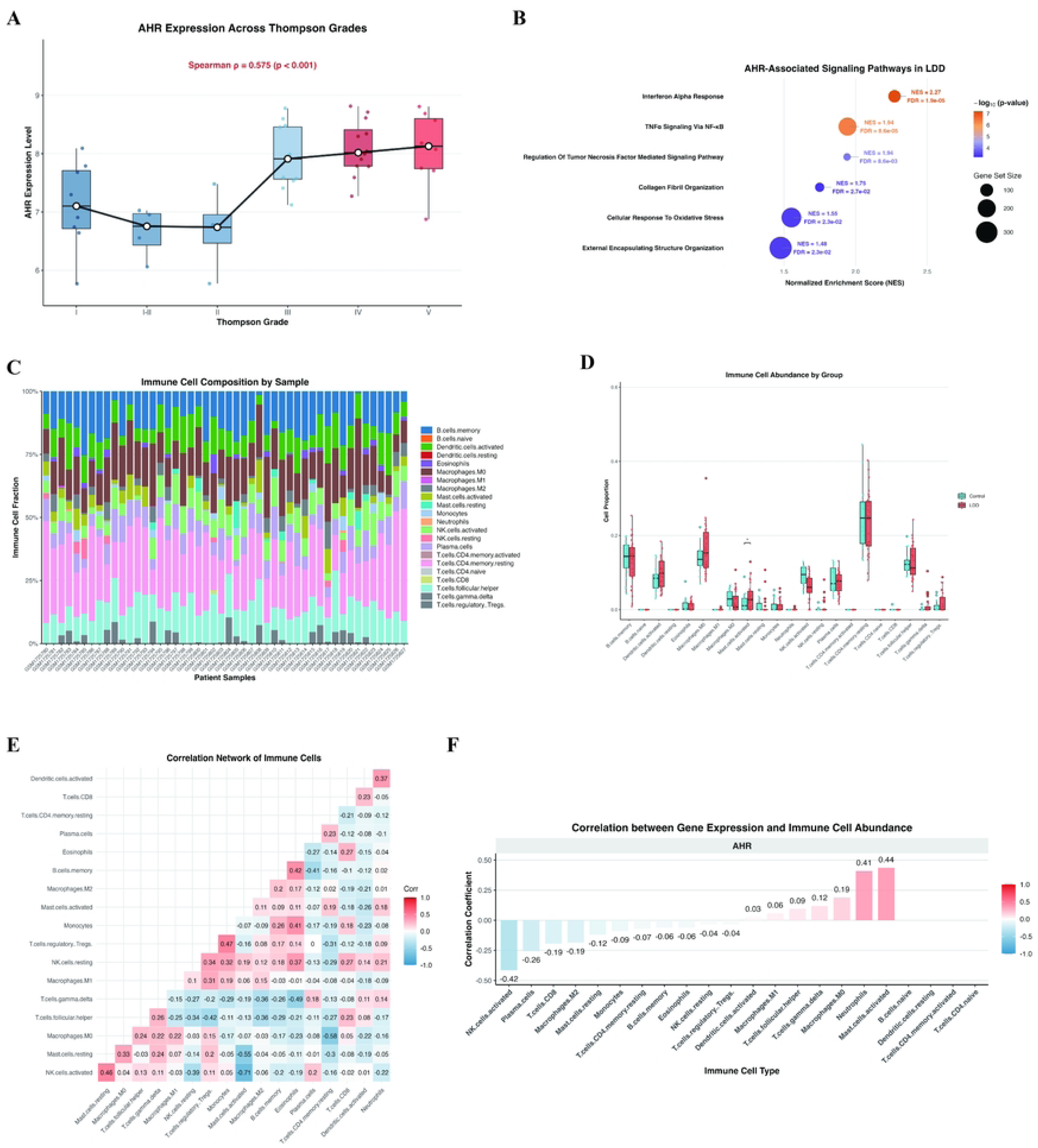
AHR expression dynamics and immune infiltration in LDD. (A) AHR expression across Thompson grades, showing progressive upregulation with increasing degeneration severity. (B) GSEA highlighting AHR-associated signaling pathways. (C) Stacked bar plot showing relative immune cell composition in normal versus degenerative disc tissues. (D) Boxplots depicting differential abundance of immune cell subsets, with activated mast cells significantly enriched in LDD. (E) Correlation matrix of immune cell subsets, illustrating inter-relationships across the immune microenvironment. (F) Correlation between AHR expression and immune cell abundance, demonstrating a positive association with activated mast cells and other inflammatory subsets.

### 3.12 AHR-Related Pathways Identified by GSEA

To further investigate the potential role of AHR in intervertebral disc degeneration, we performed single-gene GSEA analysis based on the GSE70362 dataset (Figure 7B). The analysis demonstrated that AHR-associated genes were significantly enriched in inflammation- and immune-related pathways, including interferon-α response, TNFα signaling via NF-κB, and TNF-mediated signaling. In addition, enrichment was observed in the cellular response to oxidative stress, as well as in structural processes such as collagen fibril organization and ECM organization. These findings suggest that AHR is closely linked to inflammatory activation, oxidative stress, and ECM remodeling in intervertebral disc degeneration.

### 3.13 Immune Landscape Characterization in LDD

Immune infiltration analysis delineated the immune-cell composition across samples, with most subsets showing comparable proportions between control and degenerative discs (Figure 7C). Among all examined populations, only activated mast cells displayed a statistically significant increase in the degenerative group, whereas the remaining subsets were nonsignificant (Figure 7D). The immune-cell correlation network suggested coordinated fluctuations among subsets without evidence of a global immune shift (Figure 7E). AHR expression correlated positively with activated mast cells, while correlations with other cell types were weak or negligible, reinforcing a selective association between AHR activity and mast-cell activation in IDD (Figure 7F).

### 3.14 PCR and Western blot validation of core targets

Quantitative PCR analysis confirmed that AHR mRNA expression was significantly elevated in the LDD group compared with controls, whereas treatment with Xiaobi Plaster (LDD+XBP) partially reduced this increase (Figure 8A). Similar but less pronounced trends were observed for PTPN2 and SOAT1. At the protein level, Western blot assays further validated these findings (Figure 8B–C). The inclusion of a sham-operated group confirmed that puncture injury itself induced an increase in AHR, PTPN2, and SOAT1 protein levels. XBP administration attenuated, but did not fully normalize, these elevations. Among the three targets, AHR showed the most prominent dysregulation and the clearest response to treatment, while PTPN2 and SOAT1 displayed more moderate changes.

**Figure 8.**
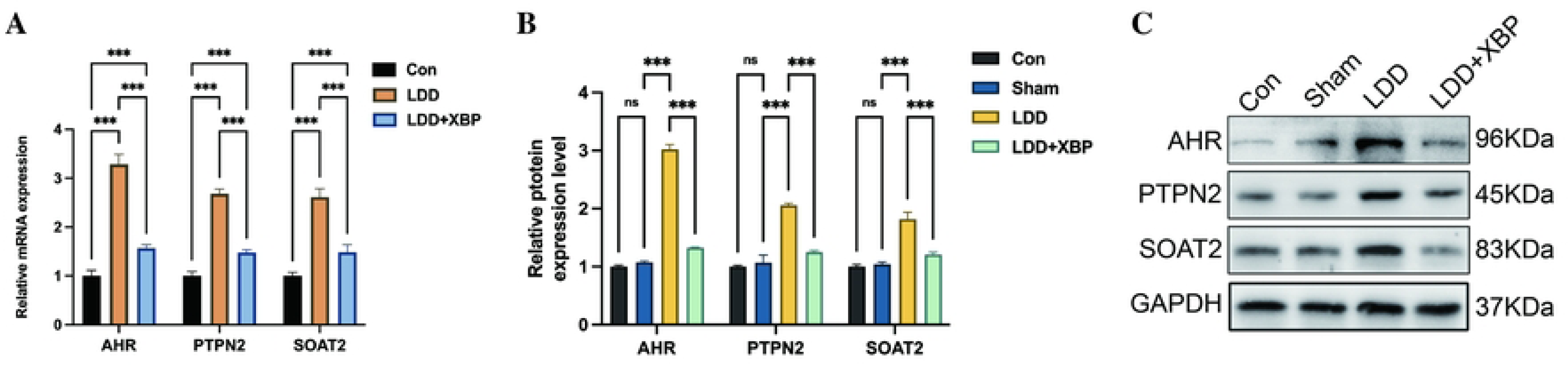
Validation of candidate targets in rat LDD models. (A) qPCR analysis showed that AHR, PTPN2, and SOAT1 mRNA expression was significantly upregulated in LDD compared with controls, while Xiaobi Plaster (XBP) treatment partially reversed these changes. (B) Relative protein expression quantified from Western blot assays showed a similar trend. (C) Representative Western blot images of AHR (96 kDa), PTPN2 (45 kDa), and SOAT1 (83 kDa), with GAPDH (37 kDa) as loading control. Data are shown as mean ± SD; ns, not significant; ***p < 0.001.

## 4 Discussion

The present study provides novel mechanistic insights into how XBG may contribute to the treatment of LDD. By integrating network pharmacology, transcriptome-driven enrichment analysis, and molecular docking/dynamics with immune profiling and limited experimental validation, we identified multiple bioactive compounds and diverse signaling pathways potentially involved in disc homeostasis. Among the predicted targets, AHR emerged as a central regulator, showing stable binding with rutaecarpine and strong association with inflammation-, ECM-, and immune-related processes. These findings suggest that XBG acts through a multi-target, multi-pathway mode of action, in which the AHR signaling axis represents a pivotal hub linking immune and extracellular matrix remodeling in LDD.

AHR is a ligand-activated transcription factor and a pioneer member of the basic helix–loop–helix/Per-ARNT-Sim family, functioning as a cellular sensor for both exogenous and endogenous signals [51,52]. In response to such ligands, AHR translocates to the nucleus and regulates a wide gene repertoire including those encoding cytochrome P450 enzymes, orchestrating xenobiotic metabolism and detoxification [52]. Beyond its canonical detoxifying role, AHR is increasingly recognized as a key regulator of inflammation, oxidative stress responses, immune cell differentiation, and ECM remodeling—processes central to the development of LDD [52,53]. Given that LDD involves chronic inflammatory signaling, oxidative damage, and matrix degradation, AHR’s multifaceted regulatory functions align closely with these pathological hallmarks. Moreover, AHR is highly expressed in barrier tissues like the skin, where it governs keratinocyte differentiation, cutaneous immune responses, and oxidative defense mechanisms—traits particularly relevant given our use of a transdermal herbal formulation in this study[54]. This abundance of AHR in the skin suggests that rutaecarpine may activate cutaneous AHR upon topical administration, thereby triggering systemic anti-inflammatory or immunomodulatory pathways that ultimately affect disc tissue integrity. Such a dual role—not only as a core mediator in disc pathology but also as a key interface for transdermal drug action—cements AHR’s relevance both biologically and therapeutically in our study.

In addition to AHR, two other validated targets—PTPN2 and SOAT1—emerged in our analyses, potentially contributing to LDD through distinct yet synergistic mechanisms. PTPN2, a non-receptor protein tyrosine phosphatase, is widely recognized as a crucial negative regulator of inflammation. It dampens inflammatory cascades by dephosphorylating key signaling molecules such as STAT3 and p65, thereby limiting macrophage activation and cytokine production [55]. Notably, PTPN2 deficiency has been shown to exacerbate inflammatory responses and disrupt barrier integrity in epithelial models [56]. In the context of LDD, PTPN2 may serve as a protective brake against uncontrolled inflammatory signaling and oxidative injury.

Meanwhile, SOAT1 (sterol O-acyltransferase 1) catalyzes the esterification of free cholesterol into cholesterol esters, thereby maintaining intracellular lipid homeostasis. In pathological conditions such as hepatocellular carcinoma, SOAT1 overexpression promotes epithelial–mesenchymal transition (EMT) and enhances invasive potential by increasing cholesterol esterification, highlighting the link between lipid metabolic dysregulation and ECM remodeling [57]. Although direct data in intervertebral disc degeneration are not yet available, such mechanisms suggest that elevated SOAT1 activity may similarly disrupt membrane lipid composition, impair disc cell homeostasis, and exacerbate biomechanical vulnerability of the ECM. Thus, SOAT1 may serve as a metabolic modulator that intersects lipid dysfunction with structural degeneration in LDD. The dysregulation of AHR, PTPN2, and SOAT1 delineates a multi-layered regulatory axis in LDD, integrating inflammatory signaling, oxidative– metabolic balance, and ECM remodeling as interconnected determinants of disc homeostasis.

In the GSE70362 dataset, differential expression analysis followed by GO and KEGG enrichment revealed a landscape dominated by pathways related to inflammation, immune responses, oxidative stress imbalance, and ECM remodeling. Significantly enriched processes included the TNF/NF-κB axis, IL-17 signaling, cytokine–receptor interactions, MAPK signaling, ECM–receptor interaction, and focal adhesion. These pathways are highly consistent with the established pathophysiology of LDD, which is characterized by chronic low-grade inflammation, ROS-driven injury, and progressive ECM degradation[2,3,58,59]. Building upon this enrichment landscape, we then performed a single-gene GSEA centered on AHR. Notably, genes co-expressed with AHR were significantly enriched in pathways closely mirroring those identified in the DEG analysis, including interferon-α response, TNFα/NF-κB signaling, and oxidative stress responses, as well as ECM structural organization and collagen fiber assembly. This overlap underscores that AHR is not merely one of many dysregulated genes, but rather a regulatory hub integrating inflammatory and redox signals with ECM remodeling. Mechanistically, AHR exhibits bidirectional crosstalk with NF-κB signaling, and ligand activation of AHR can attenuate p65/RelA activity and inflammatory cytokine output in specific contexts; in parallel, AHR signaling is linked to extracellular-matrix remodeling through the regulation of matrix metalloproteinases, supporting its role in ECM turnover [60–63]. Such evidence supports the notion that AHR may serve as a potential upstream coordinator linking immune activation and tissue remodeling in the degenerative disc microenvironment. Taken together, the convergence between DEG-level enrichment and AHR-specific enrichment reinforces AHR as a promising therapeutic target in LDD.

Among the candidate active molecules predicted from the network pharmacology– based compound–target screening, rutaecarpine emerged as the most stable ligand for AHR, as supported by both molecular docking and molecular dynamics simulations. Rutaecarpine has been shown to activate Nrf2/HO-1 antioxidant defenses and suppress NF-κB–mediated inflammatory signaling in gastric injury models, supporting its dual anti-inflammatory and antioxidant effects [64]. Additionally, rutaecarpine inhibits the KEAP1–Nrf2 interaction, further facilitating Nrf2 activation in models of colitis [65]. These mechanisms are directly relevant to LDD, which is marked by chronic inflammation, oxidative stress, and ECM degradation [66,67]. AHR is known to mediate matrix remodeling—e.g., its activation induces MMP-1 production, a key factor in ECM turnover [68]. Also, AHR deficiency is associated with dysregulated matrix metabolism in age-related tissue degeneration [69]. Together, these findings support a coherent mechanistic link whereby rutaecarpine, through selective engagement of AHR, may attenuate inflammatory and oxidative stress responses while simultaneously modulating ECM turnover.

Immune deconvolution revealed a largely unchanged leukocyte landscape between normal and degenerative discs, with one consistent exception: activated mast cells were significantly enriched in LDD. This pattern aligns with tissue studies showing that mast cells are present in painful, degenerate human discs and that mast cell–disc cell crosstalk amplifies inflammatory, catabolic, and pro-angiogenic programs implicated in degeneration and pain [70,71]. Notably, AHR expression positively correlated with mast-cell infiltration, a relationship that is biologically plausible because mast cells express functional AHR and AHR ligands can modulate mast-cell differentiation and activation via Ca²⁺/ROS-dependent signaling [72]. Endogenous tryptophan metabolites such as kynurenine can also activate mast cells through AHR, further linking AHR tone to mast-cell activity within inflamed tissues [73]. Together, these results highlight activated mast cells as a selectively perturbed immune subset in LDD and support a mechanistic axis connecting AHR signaling and mast-cell–driven inflammation in the degenerative disc microenvironment [74].

This study has several limitations that should be acknowledged. First, our analyses relied primarily on transcriptomic datasets and computational predictions, which may not fully reflect the cellular complexity of degenerative disc tissues. Incorporating single-cell or proteomic data could provide more refined insights into the cell type– specific roles of AHR, PTPN2, and SOAT1. Second, while docking and molecular dynamics simulations identified rutaecarpine as a stable AHR ligand, experimental assays are needed to confirm this interaction. Similarly, the association between AHR and mast cell infiltration was correlative; functional studies, such as in vitro co-culture or in vivo gene manipulation, are required to establish causality. Third, the puncture-induced rat model provided supportive evidence but cannot fully replicate the chronic and multifactorial nature of human LDD. Larger-scale patient-based studies and integrative multi-omics approaches will be essential to validate translational relevance. Finally, although rutaecarpine emerged as a promising candidate, Xiaobi Plaster is a multi-component formulation. Future research should consider potential synergistic or antagonistic effects among its active compounds.Addressing these limitations will help refine our understanding of AHR biology and accelerate its translation into a therapeutic target for LDD.

## 5 Conclusion

This study integrated network pharmacology, machine learning, molecular docking and dynamics, enrichment analyses, and experimental validation to elucidate the mechanisms of XBG in LDD. We identified AHR as a central regulatory hub, with stable binding to rutaecarpine and enrichment in inflammation, oxidative stress, and ECM remodeling pathways. PTPN2 and SOAT1 were also implicated as auxiliary modulators of inflammatory and metabolic balance. Transcriptomic validation confirmed AHR expression correlated with degeneration grade and mast cell infiltration. These findings highlight a multi-target mode of action for XBG and support AHR as a promising therapeutic target in LDD.

## Data availability statement

The datasets analyzed in this study are publicly available and were not generated by the authors. Specifically, the transcriptomic datasets GSE70362 and GSE245147 were retrieved from the NCBI Gene Expression Omnibus (GEO) database (http://www.ncbi.nlm.nih.gov/geo/). All bioinformatic analyses were conducted using these publicly accessible resources. Additional data supporting the findings of this study are available from the corresponding author upon reasonable request.

## Ethics Statement

All animal experimental procedures were conducted in strict accordance with the Guidelines for the Care and Use of Laboratory Animals issued by the National Institutes of Health (NIH, USA). The study protocol was reviewed and approved by the Experimental Animal Ethics Committee of the First Affiliated Hospital of the University of Science and Technology of China (Approval No.: 2023-N(A)-183).

Efforts were made to minimize the number of animals used and to reduce their suffering.

## Author Contributions

Huaize Wang, Jiejie Sun, Bin Dai, Jiafeng Peng, Hongxing Zhang, Zhiwen Zheng, Minglei Gao, Ran Xu, Junchen Zhu, and Yingzong Xiong contributed to this work. Huaize Wang performed animal experiments and molecular validations; Jiejie Sun conducted immune infiltration analysis and assisted in data interpretation; Bin Dai supported clinical data interpretation and manuscript revision; Jiafeng Peng and Hongxing Zhang carried out bioinformatics analyses, molecular docking, and manuscript drafting; Zhiwen Zheng, Minglei Gao, and Ran Xu contributed to data curation, visualization, and literature review; Yingzong Xiong provided technical assistance; and Junchen Zhu conceived and supervised the study, coordinated the project, and finalized the manuscript. All authors read and approved the final version of the manuscript.

## Supplementary Figure

**Supplementary Figure 1** Concordance of DEG expression between NP and AF. Correlation of gene expression changes between nucleus pulposus (NP) and annulus fibrosus (AF). Log2 fold changes from NP and AF showed strong concordance (Spearman ρ = 0.908; Pearson r = 0.905), supporting the reliability of the combined DEG analysis.

## Supplement Table

**Supplementary Table S1.** Compound–target dataset for Xiaobi Plaster. All herbal components, active compounds, and predicted targets were collected from TCMSP and SwissTargetPrediction databases for subsequent network pharmacology analysis.

**Supplementary Table S2.** Primer sequences used for qRT-PCR validation of AHR. Forward and reverse primers for AHR are shown.

